# Proximity and Touch are associated with Neural but not Physiological Synchrony in Naturalistic Mother-Infant Interactions

**DOI:** 10.1101/2021.01.21.427664

**Authors:** Trinh Nguyen, Drew H. Abney, Dina Salamander, Bennett I. Bertenthal, Stefanie Hoehl

## Abstract

Caregiver touch plays a vital role in infants’ growth and development, but its role as a communicative signal in human parent-infant interactions is surprisingly poorly understood. Here, we assessed whether touch and proximity in caregiver-infant dyads are related to neural and physiological synchrony. We simultaneously measured brain activity and respiratory sinus arrhythmia of 4- to 6-month-old infants and their mothers (N=69 dyads) in distal and proximal joint watching conditions as well as in an interactive face-to-face condition. Neural synchrony was higher during the proximal than during the distal joint watching conditions, and even higher during the face-to-face interaction. Physiological synchrony was highest during the face-to-face interaction and lower in both joint watching conditions, irrespective of proximity. Maternal affectionate touch during the face-to-face interaction was positively related to neural but not physiological synchrony. This is the first evidence that touch mediates mutual attunement of brain activities, but not cardio-respiratory rhythms in caregiver-infant dyads during naturalistic interactions. Our results also suggest that neural synchrony serves as a biological pathway of how social touch plays into infant development and how this pathway could be utilized to support infant learning and social bonding.

**Highlights:**

- Mother-infant neural and physiological synchrony arise during mutual engagement.
- Behavioral correlates of neural and physiological synchrony diverge.
- Interpersonal neural synchrony is related to proximity and affective touch.
- Interpersonal physiological synchrony is related to infant negative affect.

## 1. Introduction

Human development is driven by social interactions. From early on, infants begin to actively seek information as embodied agents in their social interactions (De Jaegher et al., 2016; Raz and Saxe, 2020). These exchanges are essential for infants’ developing understanding of self and others. In these interactions infant and caregiver typically fluctuate between aligned and misaligned states (Montirosso and McGlone, 2020). Communicative signals allow the dyad to “repair” such misaligned states (Tronick & Gianino, 1986). This kind of coordination is referred to as interpersonal synchrony (Feldman, 2012) or mutual attunement (Stern et al., 1985), a dynamic process by which behavior and neurophysiological processes are reciprocally adjusted between two or more persons. Temporally aligning with another person facilitates mutual prediction and allostasis, i.e., the ongoing interpersonal physiological regulation required to meet the changing demands of the environment (Atzil et al., 2018). The occurrence of early interpersonal synchrony in various modalities is evidenced at 3-4 months of age, when the social capacities of infants begin to emerge (Beebe et al., 2010; Feldman, 2012). Infants begin to perceive contingent relations, discriminate the strength of these relations, and generate predictions based on these contingencies (e.g., Beebe et al., 2016; Harrist & Waugh, 2002). By this age, caregiver and infant have had a sufficient amount of interaction history, so the infant’s predictions of the interactive time course with their caregiver become more precise, highlighting the infant’s communicative capacity (Tronick, 1989). Interpersonal synchrony occurs on the behavioral, neural and physiological level, yet we know little about their relational and developmental dynamics (Beauchaine, 2015; Feldman, 2017). Here, we simultaneously measured the brain activity and physiology of 4- to 6-month-old infants and their mothers during non-interactive and interactive contexts utilizing a multi-method “hyperscanning” setup to investigate the role of social touch, as a communicative signal, on caregiver-infant neural and physiological synchrony.

Recent research with adults using simultaneous recordings of brain activities from several persons (hyperscanning) demonstrates that interactive synchrony also manifests as interpersonal synchronization of brain activities (Dumas et al., 2011; Hasson et al., 2012). This interpersonal neural synchrony (INS) is thought to facilitate turn-taking interactions (Wilson and Wilson, 2005) and interpersonal transmission of information through verbal and non-verbal communication (Dumas et al., 2011; Hasson et al., 2012). A facilitative function of touch for INS was highlighted by a recent study showing that handholding in adults increases INS (Goldstein et al., 2018). INS has been mainly identified in brain regions associated with socio-cognitive processes (Gvirts and Perlmutter, 2020; Hoehl et al., 2020; Koban et al., 2019; Redcay and Schilbach, 2019), including mutual attention, prediction (prefrontal cortex), affect sharing (inferior frontal gyrus), mentalizing and shared intentions (temporo-parietal junction).

By contrast, the neural mechanisms of social exchanges in early child development are still poorly understood as few developmental hyperscanning studies exist to date. The emerging evidence demonstrates that adults and children synchronize their brain activities to a greater degree in interactive contexts that require mutual engagement than when solving the same tasks individually or when interacting with a third person (Nguyen et al., 2020, 2021; Piazza et al., 2020). Importantly, behavioral indicators of high mutual attunement, such as reciprocity and eye contact were linked with heightened levels of INS. These studies suggest that INS could be a sensitive biomarker for successful attunement between caregivers and their infants. However, social touch has not yet been considered in caregiver-infant neural synchronization.

Synchronous interactions on the behavioral level have also been linked to physiological synchronization (IPS). Respiratory sinus arrhythmia (RSA) is an index of the functioning of the vagal system. The regulation of the vagal system allows us to adapt to our ever-changing social environments (Porges, 2007). The attunement of cardio-respiratory rhythms between mother and child emerges very early in life and is suggested to function as a scaffold for infants’ still immature physiological systems (Abney et al., 2021a; Feldman et al., 2011). The coupling of physiological states thereby supports the development of infants’ self-regulation and various other neurobehavioral and physiological functions (Atzil et al., 2018). Overall, the importance of infant-parent RSA co-regulation for properties and outcomes of social interaction like emotion regulation and distress provides the basis for why we focus on the physiological measure in the current study.

Research on early IPS underscores that physiological alignment occurs under conditions of mutual engagement and decreases when interacting partners disengage (Feldman et al., 2011). More specifically, IPS seems to emerge after instances of infants showing high arousal and negative affect in daily life (Wass et al., 2019). After a negative affect manipulation, 12- to 14-month-old infants displayed IPS with their mothers when the infant was seated on the mother’s lap in comparison to a no-touch condition (Waters et al., 2017). Though preliminary, findings speak to a potential role of touch in affect contagion through physiology. Across development, there is growing evidence pointing towards the role of touch in IPS (Goldstein et al., 2017; Van Puyvelde et al., 2015). Yet, a systematic investigation of the role of spontaneous maternal touch on IPS during a naturalistic interaction with the infant is still lacking.

This study integrates the concurrent assessment of INS and IPS in a multi-level hyperscanning paradigm with mother-infant dyads naturally interacting with each other. We tested whether attunement through proximity and touch between the dyad is associated with INS and IPS. INS was assessed in 4- to 6-month-old infants and their primary caregivers by using dual-functional near-infrared spectroscopy (dual-fNIRS). As aforementioned, social interactions at this age provide a unique opportunity to study infants’ social capacities to synchronize with their caregivers (Beebe et al., 2010). Importantly, we limited the age range to 6 months of age, as by then the primacy of face-to-face exchanges tends to decrease as infants begin to increasingly explore physical objects (Jaffe et al., 2001). We focused on prefrontal and inferior frontal regions. Synchronization in these regions is suggested to mark mutual attention and shared affect, which are critical processes to infants’ interactions with their caregivers (Feldman, 2017). We further assessed IPS through electrocardiography (ECG) and coded the dyad’s behavior from video recordings.

According to our pre-registered hypotheses, we expected caregiver-infant dyads to show increased INS and IPS during phases of high, experimentally manipulated physical proximity between mother and infant. We contrasted the dyads’ INS and IPS during a distal joint watching condition to a proximal joint watching condition while caregiver and infant were attending to the same video. Subsequently, we assessed whether spontaneous use of maternal touch during a face-to-face interaction was related to differences in INS and IPS. We hypothesized that increased durations of social touch would be associated with increased INS and IPS. Additionally, we studied infants’ affect in relation to INS and IPS and examined potential links between INS and IPS in the mother-infant dyad.

## 2. Material and Methods

### 2.1 Participants

Overall, 81 mother-infant dyads participated in the present study and were recruited from a database of volunteers. Out of those dyads, 72 completed the experiment, while the procedure was incomplete for 9 dyads due to fussiness (infants started to cry during the preparation phase or before the end of the experiment), and sleepiness. Due to the novel paradigm and lack of effect sizes, we aimed for a final sample size of 50 infants and oversampled for expected attrition rates (∼30%). Infants’ age ranged from 4- to 6-months-old (*M*=4.7 months; *SD*=16 days; 33 girls). Infants were born healthy and at term, with a gestation period of at least 36 weeks. Mothers’ age averaged 33.97 years (*SD*=4.94) and 57% of mothers had a university degree. All dyads were of White European origin and came from middle to upper-class families based on parental education. All infants and mothers had no neurological problems as assessed by maternal reports. The study was approved by the local ethics committee. Parents provided written informed consent on behalf of their infants and themselves. Participation was remunerated.

### 2.2 Experimental procedure

During the experiment, caregiver and infant were either seated next to one another or the infant sat on the caregiver’s lap while watching a calm aquarium video on a tablet (distal watching and proximal watching conditions; see *Figure 1*). The videos lasted 90 sec. and depicted fish swimming in a tank. The order of the watching conditions was counterbalanced. Next, mother and infant engaged for 5 min. in free play without toys and song while both were seated face-to-face (interactive free play condition). Neural activity in the mother-infant dyad was simultaneously measured with fNIRS. We assessed RSA through ECG and each dyad was filmed by three cameras (angled towards the dyad, the infant, and the mother) throughout the experiment.

**Figure 1.**
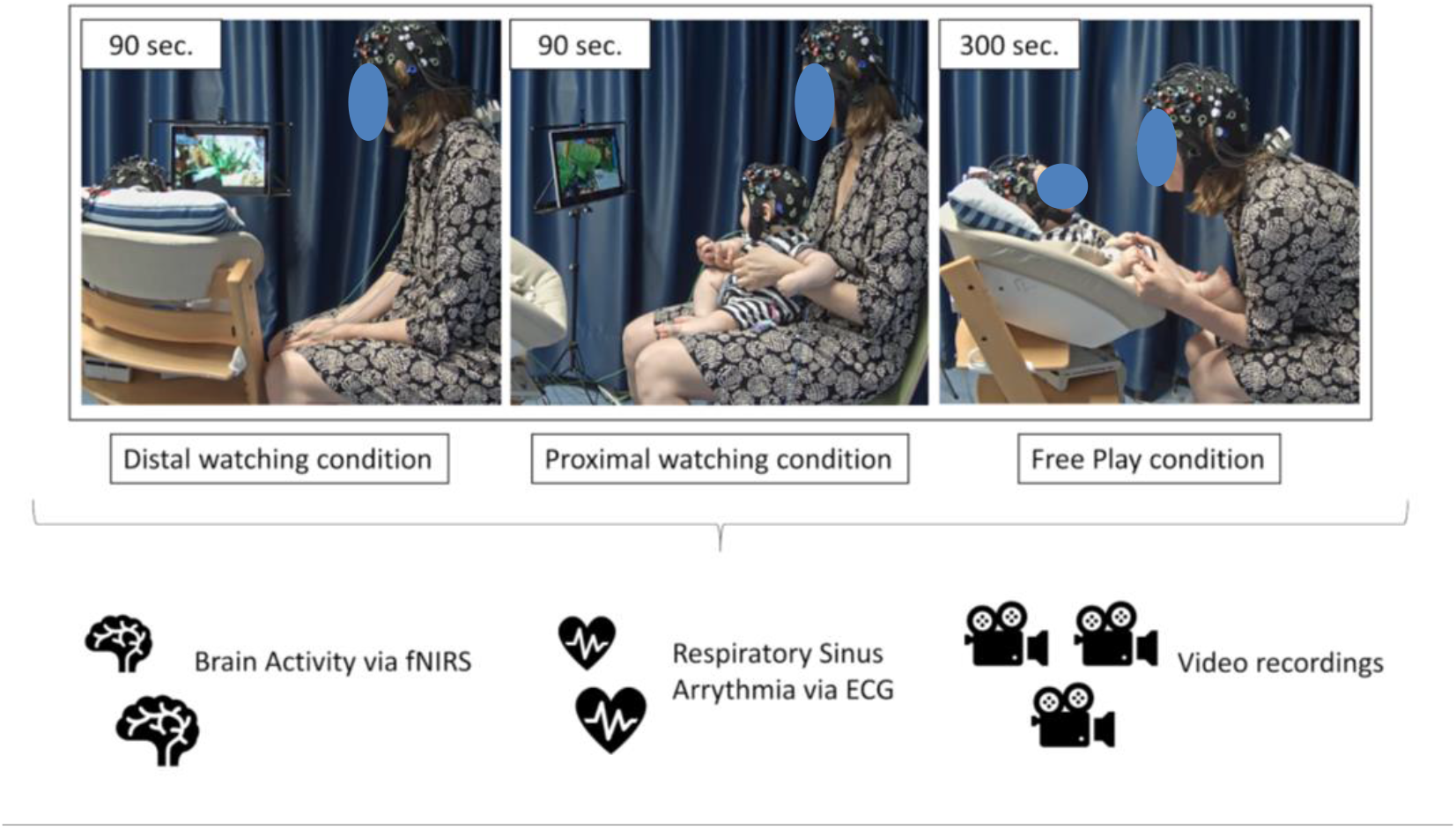
An exemplary mother-infant dyad during the joint watching conditions with and without physical contact (90 seconds) and the free play interaction condition (300 seconds) (from left to right). Throughout the experiment, we simultaneously measured brain activity via functional near-infrared spectroscopy (fNIRS), respiratory sinus arrhythmia (RSA) via electrocardiography (ECG) and subsequently coded the dyads behavior through video recordings. The three cameras were facing the infant, the mother, and the dyad, respectively.

### 2.3 Data acquisition and processing

#### 2.3.1 fNIRS Recordings

We used two NIRSport 8-8 (NIRx Medizintechnik GmbH, Germany) devices to simultaneously record oxy-hemoglobin (HbO) and deoxy-hemoglobin (HbR) concentration changes in mother and infant. The 8 × 2 probe sets were attached to an EEG cap with a 10-20 configuration (*Figure S1*). The probe sets over the left and right inferior frontal gyrus (IFG) surrounded F7 and F8, whereas the probes on the medial prefrontal area (mPFC) surrounded FP1 and FP2. These regions of interest were based on previous work involving adult-child interactions (Nguyen et al., 2020; Piazza et al., 2020; Redcay and Schilbach, 2019). In each probe set, 8 sources and 8 detectors were positioned, which resulted in 22 measurement channels with equal distances of ∼2.3 cm between the infants’ optodes and 3 cm between the mothers’ optodes. The absorption of near-infrared light was measured at the wavelengths of 760 and 850 mm and the sampling frequency was 7.81 Hz.

fNIRS measurements were processed using MATLAB-based functions derived from Homer 2 (Huppert et al., 2009). Raw data was converted into optical density. Next, optical density data were motion-corrected with a wavelet-based algorithm with an interquartile range of 0.5. Motion-corrected time series were further visually inspected during a quality check procedure. Before continuing, 22.87% of the channels from both mother and child were removed from further analyses due to bad signal-to-noise ratio and motion artifacts. Then slow drifts and physiological noise were removed from the signals using a band-pass second-order Butterworth filter with cutoffs of 0.01 and 0.5 Hz. The filtered data were converted to changes (μMol) in oxygenated (HbO) and deoxygenated hemoglobin (HbR) based on the modified Beer-Lambert Law. For later analyses, both HbO and HbR synchrony are reported.

#### 2.3.2 Electrocardiography (ECG) Recordings

We made use of a Brain-Amp system (Brain Products GmbH, Germany) with two amplifiers to measure two standard single-channel ECG registrations (lead II derivation). One electrode was placed on the upper right chest, one on the left side of the abdomen, and the grounding electrode was placed on the right side of the abdomen on both infant and mother. The ECG signal was recorded with a 500 Hz sampling frequency.

Interbeat-intervals (IBIs) were extracted offline using ARTiiFACT (Kaufmann et al., 2011). The ECG data were visually inspected for (in)correct detections and artifacts by trained research assistants. When ectopic beats or erroneous detections were found, the data were manually corrected (removal of erroneous detection/artifact followed by a cubic spline interpolation; corrections < 1%). Next, IBIs were down-sampled to 5 Hz and a 51-point band-pass local cubic filter was used to estimate and remove the slow periodic and aperiodic components of the time-series. A FIRtype bandpass filter was applied to further isolate the variance in the IBI series to only the frequency range of spontaneous breathing for infants (0.3-1.3 Hz) and adults (0.12-1.0 Hz). The range for mothers’ respiration was expanded from its typical value of 0.4 to 1.0 to account for the infrequent occurrence of faster breathing during talking or playing segments so that the same filter could be used for all mothers in all conditions. The Porges & Bohrer (1990) technique for RSA magnitude estimation includes parsing this component signal into discrete epochs (lasting 10 to 120 sec), then calculating the natural log of the variance in each epoch. RSA is reported in units of ln(ms)^2^. In order to collect a more continuous measure of RSA, a sliding window of 15 seconds was used to extract a continuous (updated every 200 ms) estimate of cardiac vagal tone for both participants. The estimated RSA value corresponded to the first value of the sliding window (see Abney et al., 2021b for detailed information).

#### 2.3.3 Synchrony estimation

We used Wavelet Transform Coherence (WTC) and Cross-Recurrence Quantification Analyses (CRQA) to estimate INS and IPS, respectively. WTC and CRQA are non-linear approaches to estimate synchrony in two non-stationary time-series, as we cannot assume stationarity for HbO, HbR and RSA time-series. Both methods allow the quantification of dynamical systems and their trajectories. Utilizing these methods, we are able to capture many properties of the neural and physiological dynamics that would otherwise be lost due to averaging with more traditional correlation analysis.

We assessed the relation between the fNIRS time series in each caregiver and infant using Morlet WTC as a function of frequency and time (Grinsted et al., 2004). WTC is more suitable in comparison to correlational approaches, as it is invariant to interregional differences in the hemodynamic response function (HRF) (Sun et al., 2004). Correlations, on the other hand, are sensitive to the shape of the HRF, which is assumed to be different between individuals (especially of different ages) as well as different brain areas. Moreover, a high correlation may be observed among regions that have no blood flow fluctuations. In previous hyperscanning research in both adults and children, the analyses have shown that synchronization occurs in different frequency bands (e.g., Cui et al., 2012; Jiang et al., 2012; Zhao et al., 2021). For the current study, we knew little about the relevant frequency bands (especially in parent-infant interactions), and thus we determined the frequency band after we estimated the coherence between mother and infant. Accordingly, based on previous studies (Nguyen et al., 2020, 2021), visual inspection, and spectral analyses, the frequency band of 0.012 Hz – 0.312 Hz (corresponding to 8 - 32 s) was identified as the frequency-band of interest. Average coherence (INS) was then calculated for the distal watching condition, the joint watching condition, and the interactive free play condition in each channel, which resulted in 3 (conditions) x 22 (channels) coherence values for each dyad. To ensure that the different durations between the first two and last conditions did not confound coherence values, we also estimated WTC in 90 second epochs in the free play condition, and subsequently compared the values to the joint watching conditions while controlling for duration. The results are described in the Supplement S1.

To examine IPS, we used cross-recurrence quantification analysis (CRQA) (Coco and Dale, 2014) to identify coupling between mothers’ and infants’ RSA time-series. CRQA is a nonlinear method for analyzing shared dynamics between two different data series and has been applied successfully to investigate cardio-respiratory dynamics (Konvalinka et al., 2011). The method is especially suitable for RSA synchrony estimations, as RSA is an estimate derived from a specific frequency band in IBIs, this renders RSA synchrony calculations unsuitable for measures of coherence. The metric we used to evaluate the RSA time-series is %DETerminism (%DET). %DET quantifies the predictability of the time-series and is calculated as the percentage of recurrent points that form diagonal lines in a recurrence plot (i.e., which are parallel to the central diagonal). Higher determinism with the same amount of recurrence implies stronger coupling. For this analysis, the recurrence rate was fixed at 2% to be able to compare CRQA estimates across conditions. We used the function optimizeParam to estimate the parameters for radius, embedding dimension(s) and delay, which resulted in radius=0.02, emb=1 and delay=16.

We also estimated the lag-0 cross-correlation between mothers’ and infants’ fNIRS as well as detrended RSA time series. Results for cross-correlation of HbO, HbR and RSA time-series are detailed in the supplements.

### 2.4 Behavioral coding

To assess maternal touch, infants’ affect, and infants’ gaze during the watching conditions (adapted from Feldman et al., 2011), trained graduate students coded video recordings of the free-play sessions using Mangold INTERACT. The experimental sessions were filmed at 25 frames per second. Maternal touching behavior, infants’ facial affect, and gaze were micro-analyzed frame-by-frame for duration and frequency. For social touch coding, we differentiated between periods of touch and no-touch (i.e., no physical contact). Within periods of touch, segments were coded for active, passive, or functional maternal touch. Segments of active touch were subsequently divided into the two categories of affectionate and stimulating touch. For infants’ facial affect, we distinguished between positive, negative, and neutral facial expressions. For infants’ gaze during watching conditions, gaze directions were differentiated between gaze to screen, gaze to mother, gaze away (refer to Table S1 for a full description of coding categories and S3 for gaze analysis). To establish inter-rater reliability, 25% of randomly chosen videos were coded by two trained coders. Inter-rater reliability was high to excellent, namely kappa=.79 for social touch, kappa=.81 for facial affect and kappa=.92 for infants’ gaze in watching conditions.

### 2.5 Statistical Analysis

All statistical analyses were calculated in RStudio (RStudio Team, 2020). We used the function glmmTMB from the R package glmmTMB (Brooks et al., 2017). Wavelet Transform Coherence (WTC) values were entered as the dependent variable of the first Generalized Linear Mixed Model (GLMM) with condition (distal watching vs. proximate watching vs. free play) and ROI (IFG vs. lPFC vs. mPFC) as fixed factors with random slopes for each ROI and condition and random intercepts of dyads. %DET was entered into a second model with condition as a fixed factor and random slope as well as random intercepts for dyads. As both WTC and %DET values are bound by 0 and 1, we assumed a beta distribution in each model. To further examine significant effects, contrasts of factors were conducted by using post-hoc analyses (emmeans) with Tukey’s Honest Significant Difference to correct for multiple comparisons. All continuous predictor variables were z-standardized, and distributions of residuals were visually inspected for each model. Models were estimated using Maximum Likelihood. Model fit was compared using a Chi-Square Test (likelihood ratio test; (Dobson, 2002).

To test for spurious correlations in both neural and physiological synchrony, we conducted bootstrapped random pair analyses (Nguyen et al., 2020; Piazza et al., 2020). Mother’s original data was randomly paired with infant data out of the sample for 1,000 permutations. WTC and %DET were calculated for each random pair. Subsequently, the average of coherence or correlation between random pairs was computed and compared against original pairs using GLMM and post-hoc contrasts (see S1, S2 for details).

Next, we used two different statistical approaches to assess the association between touch and INS and IPS. INS necessitated GLMM to account for the repeated measures (channels/regions) which are nested in each dyad. IPS in the free play condition, on the other hand, was one measure per dyad, therefore we used a linear regression to test the relation of IPS and touch.

## 3. Results

### 3.1 Interpersonal neural synchrony

First, we tested INS (estimated with WTC) in HbO concentration changes in the three experimental conditions: the distal joint watching condition vs. the proximal joint watching condition and the interactive free play condition. WTC was entered as the response variable in the GLMM, while condition and region of interest were entered as fixed and interaction effects. We assumed a random slope for all fixed and interaction effects with random intercepts for each dyad (N=69). Three dyads had no fNIRS recordings due to technical problems with the devices. The results revealed that the fixed effects for condition, χ^2^(2)=39.91, *p*<.001, and region, χ^2^(4)=33.32, *p*<.001, as well as their interaction were significant, χ^2^(8)=25.15, *p*=.001. Comparisons (using emmeans) across conditions revealed increased INS during free play and proximal watching in comparison to distal watching that were detected in bilateral lPFC and mPFC, *t*>3.10, *p*<.005. The free play condition and proximal watching condition differed in synchronization in the right lPFC and mPFC with higher INS during free play, *t*>3.06, *p*<.006. None of the conditions differed in INS in bilateral inferior frontal gyri (IFG), *p*>.071. The results are depicted in *Figure 2*. We were able to replicate the main results we found for INS in HbO with HbR and also when including equivalent epochs (90 seconds) in all three experimental conditions (see S1).

**Figure 2.**
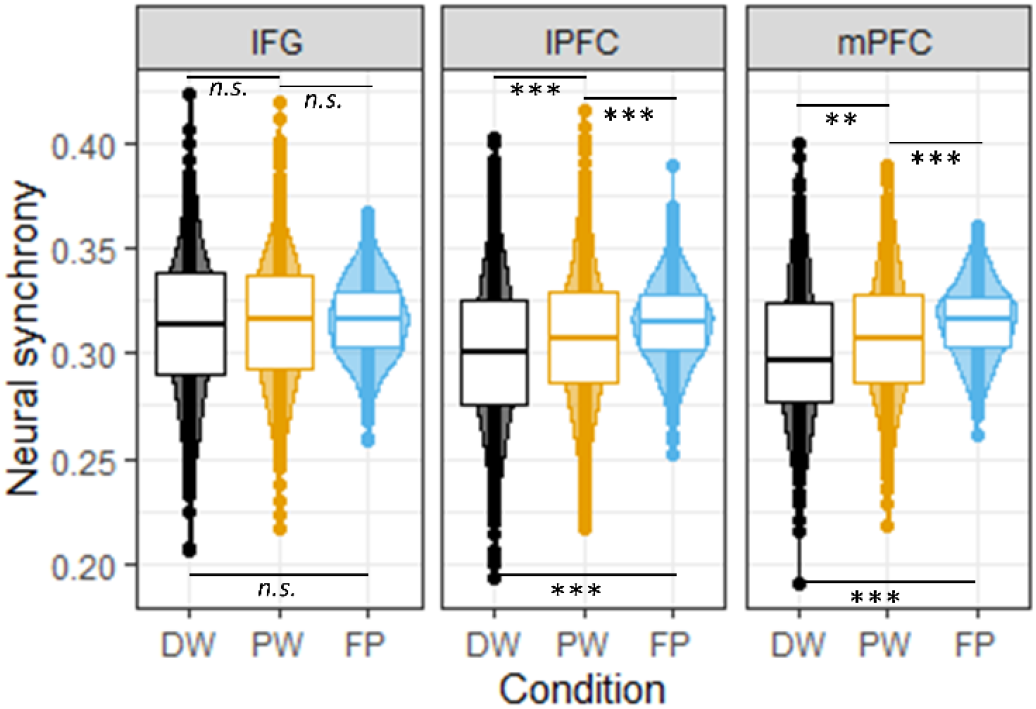
Plot of the interaction effect of condition (x-Axis) and region (facets) for INS. Neural synchrony (y-Axis) during free play was significantly higher than during distal and proximal joint watching phases in lateral and medial prefrontal areas, but not in the inferior frontal gyrus. Neural synchrony during proximate joint watching was higher than during distant joint watching, **= *p*<.010, ***= *p*<.001.

Next, we tested whether variation in social touch relates to variation in INS with a GLMM. WTC values from the free play condition were tested as the response variable. Affectionate touch, stimulating touch, passive touch, and functional touch durations were included as fixed effect variables. The results revealed a significant effect of affectionate touch, χ^2^(1)=4.13, *p*=.042 (*Figure 3A*), and stimulating touch durations, χ^2^(1)=6.24, *p*=.012 (*Figure 3B*). The model estimates show that longer durations of affectionate touch were related to higher INS, estimate=-0.079, SE=0.031, 95% CI=[0.002 0.107], whereas longer durations of stimulating touches were related to lower INS, estimate=0.055, SE=0.026, 95% CI=[-0.140 -0.018]. Fixed effects for passive touch and functional touch durations were non-significant, *p*>.122.

**Figure 3.**
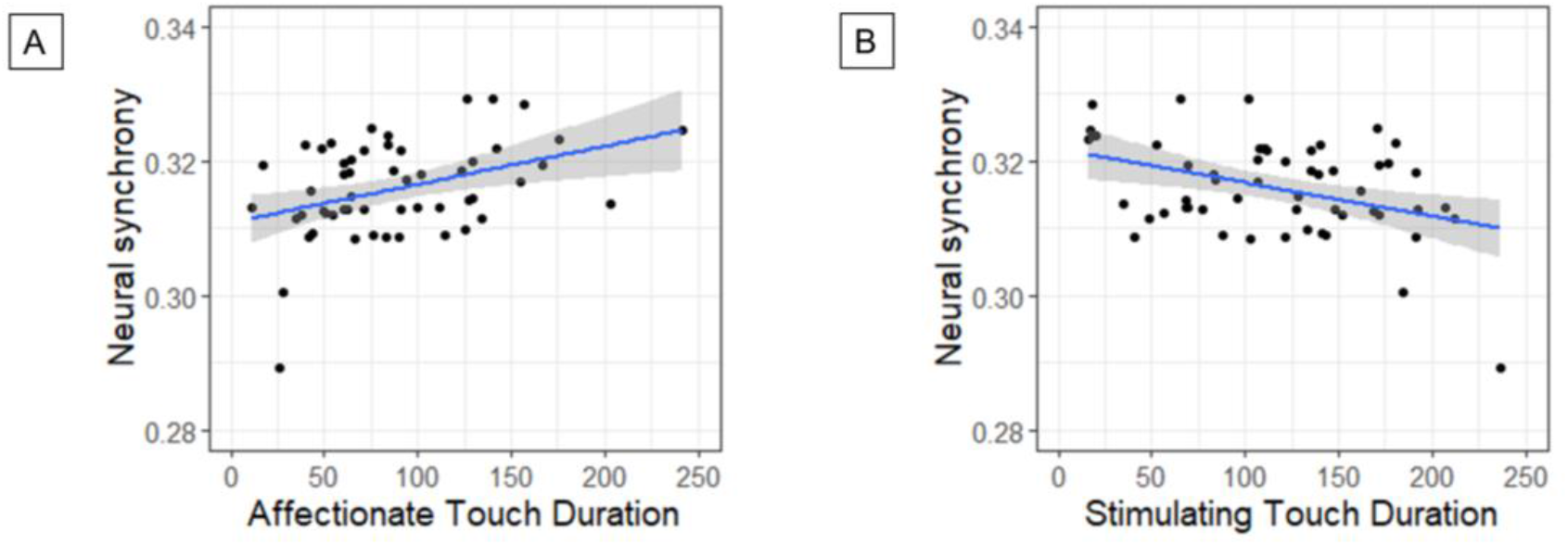
Graphs depict (A) the positive correlation between durations of affectionate touch (x-Axis) and (B) the negative correlation between durations of stimulating touch (x-Axis) and neural synchrony (y-Axis) during the free play condition. Each dot represents a dyad.

### 3.2 Interpersonal physiological synchrony

IPS between mother and infant were compared in the three experimental conditions with %DET of detrended RSA values as the response variable. Condition was entered as a fixed effect, and random intercepts for dyads were included (N=67). Five dyads were excluded due to technical problems or noisy data for which R-peaks could not be detected. The findings revealed a significant fixed effect of condition, χ^2^(2)=34.62, *p*<.001 (*Figure 4*). Post-hoc contrasts between the conditions revealed that IPS in the free play condition was significantly higher than in the distal watching condition, *t*=5.23, *p*<.001, and the proximal watching condition, *t*=4.99, *p*<.001. IPS in the joint watching conditions did not differ from one another, *p*=.978.

**Figure 4.**
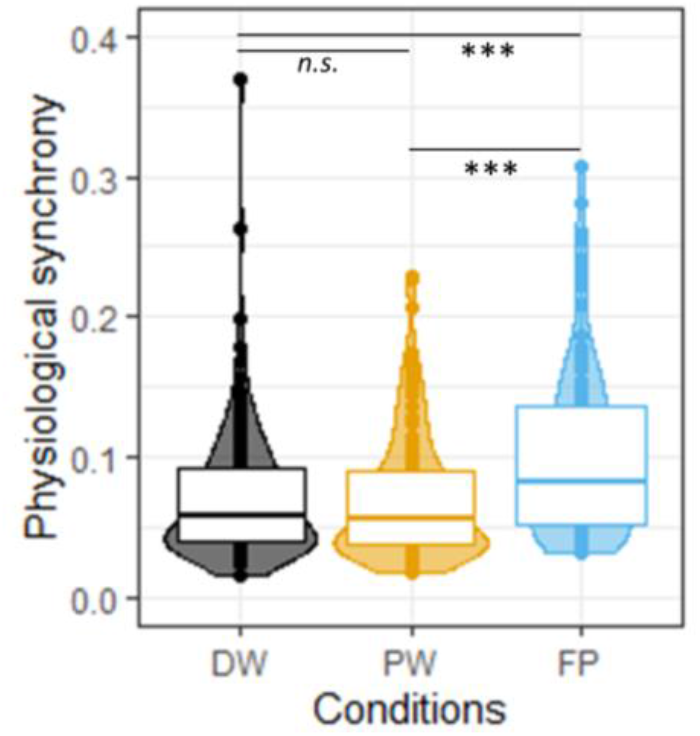
Physiological synchrony (y-Axis) during free play was significantly higher than during distal and proximal joint watching phases (x-Axis). n.s.= non-significant, **= p<.010, ***= p<.001.

Next, we tested the association between social touch and IPS with a linear regression analysis; %DET (Fisher’z-transformed) was the outcome variable. Affectionate touch, stimulating touch, passive touch and functional touch durations were included as predictor variables. While affectionate, stimulating and passive touch durations were not related to IPS, *p*>.520, functional touch durations were marginally positively associated with IPS, β=0.571, *SE*=0.237, *95% CI*=[0.091 1.051], *t*=2.48, *p*=.084.

### 3.3 Relation between INS and IPS

We assessed a potential relation between INS and IPS (both scores were averaged over each condition). HbO WTC values were entered as the response variable into the GLMM. Condition, %DET and region of interest were entered as fixed and interaction effects. We assumed a random slope for all fixed and interaction effects with random intercepts for each dyad. The results revealed no significant effect of IPS nor its interaction with conditions and/or region of interest in the brain on INS in HbO, *p*>.136.

### 3.4 Supplementary analyses

Supplementary analyses of the results showed no difference in infants’ gaze duration towards the screen between the two joint watching conditions. We also examined the role of infants’ affect to shed light on the affective context of the interaction. Overall, we find higher IPS in mother-infant dyads, but not INS, to be associated with higher infant negative affect and lower infant positive affect durations. These results are fully reported in the supplement.

## 4. Discussion

We tested the relation between mother-infant proximity and social touch and INS and IPS in three naturalistic experimental conditions with varying physical proximity and mutual engagement. Dyads displayed increased INS in bilateral lPFC and mPFC during proximity with spontaneously occurring affectionate touch. While attending to one another mother and infant showed alignment in their prefrontal brain activations, most prominently in the face-to-face interaction. Activation in the PFC is associated with the detection of communicative signals directed toward the self, mentalizing, and reward, all processes which are implicated in mutually engaged interactions (Redcay and Schilbach, 2019). Interestingly, INS in bilateral IFG did not differ between conditions and was not related to assessed behavioral correlates. IPS, on the other hand, was neither related to proximity nor spontaneous touching behavior of the mother during free play. Our findings indicate that INS in the prefrontal cortex and IPS arise in similar contexts, i.e., during mutual engagement of the mother-infant dyad, but diverge in their functionality, as they seem to be driven by different processes.

We directly contrasted the same joint watching conditions, while the proximity of mother and infant was experimentally manipulated. Joint watching of a calm video was associated with higher INS when the infant was seated on the mother’s lap instead of next to her. Physical contact allows for multi-modal stimulation and enhances caregiver responsiveness to infants (Fairhurst et al., 2014). The micro-adjustments of bodily contact and the perception of heart rhythms and respiration may have been reciprocally related to neural synchronization during this condition. Holding the infant could additionally be related to the activation of specific sensory nerve fibers through pleasant deep pressure (Case et al., 2020). We controlled for infant’s attention towards the visual stimuli (see S3), which was not different between both watching conditions. Our finding implies that proximity allows the caregiver to co-regulate their infant even when their attention is directed away from the infant, potentially preparing them for exploration or learning opportunities (Fairhurst et al., 2014).

In the free play condition, communicative signals are visibly exchanged between caregiver and child (Wass et al., 2020). Here, we identified social touch as one of the communicative signals related to INS. Specifically, affectionate touch was related to higher INS in mother-infant dyads and could serve as a parental ostensive cue (next to gaze and infant-directed speech) that entrains the infant to the ongoing pattern of early social communication (Wass et al., 2020). By eliciting infant attention, affectionate touch might help mother and infant to establish and maintain engagement with one another (Della Longa et al., 2019; Stack and Muir, 1992). The newly revealed link between affectionate touch and INS thus supports the notion that INS might be a biomarker for interaction quality. Interestingly, we find a negative association between stimulating touch and INS. These results tie in with research underlining diverging functions to different touch qualities (Mantis et al., 2019; Moreno et al., 2006). They indicate that stimulating touch, as a feature of intrusive maternal behavior, might actually be disruptive to the interaction and thus be related to lower INS (Field, 2010). Overall, the involvement of social touch in INS highlights the importance of affectionate touch in early interactions, but also the need to investigate the different qualities of touch for child development outcomes.

In addition, we simultaneously examined IPS in mother-infant dyads. IPS was, however, not related to proximity during joint watching in the current study. Previous studies also showed that mother and infant show attenuated IPS in close contact later than 3 months of age due to the widening of infants’ social orientation beyond the primary caregiver and increasing self-regulatory abilities (Van Puyvelde et al., 2015). When further probing IPS for behavioral correlates, maternal touch did not correlate with IPS between mother and infant. Instead, we find that higher IPS was associated with higher durations of infant negative affect and lower IPS was associated with higher durations of infant positive affect in the free play condition. As such, our results concur with the co-regulatory account of caregiver-infant IPS, namely its potential function to maintain allostasis (Atzil et al., 2018). The mother-infant dyads show greater physiological synchrony when co-regulation is needed, such as during infant distress marked by negative affect (Abney et al., 2021a; Wass et al., 2019). Longer durations of infants’ positive affect could have indicated that the infant was well regulated by herself. Still, more research is needed on the exact behavioral correlates of IPS to understand its functionality. For instance, considering leader-follower relations might provide further insights into IPS.

Probing the relation between INS and IPS in interactions between infants and their primary caregiver, we find both commonality and discrepancy. Passively viewing the same visual stimulation was related to lower levels of INS and IPS in comparison to a face-to-face interaction. Instead of merely reflecting common neural and physiological reactions towards the same perceptual stimulation, INS and IPS depend on the transmission of interactional signals through the environment, reflecting mutual engagement and quality of interaction (Koban et al., 2019). Though both INS and IPS were higher in the interactive than in the joint watching conditions, condition means of INS and IPS were not correlated. While INS was related to the communicative signals exchanged through touch, IPS was related to co-regulatory affective signals. This dissociation could stem from 1) the different approaches to calculate synchrony, but also suggests 2) the discrepant role of social touch for early social communication, as well as 3) a potential functional dissociation of INS and IPS. Future studies should therefore continue to combine neural and physiological assessments to dissociate the functionality of INS and IPS. Importantly, we know little about the *intraindividual* developmental trajectory of brain activation and RSA coupling (Beauchaine, 2015), which could inform *interindividual* coupling.

### 4.1 Conclusions and Future Directions

The present research is the first multi-level hyperscanning study on naturalistic mother-infant interactions with infants as young as 4 months of age. Caregiver-infant interactions between 4-6 months allow us a unique opportunity to examine infants’ developing social capacities (Feldman et al., 1996; Stern, 1985; Tronick, 1989). Importantly, these interactions robustly predict later social and cognitive development (e.g., Beebe et al., 2010; Field, 1995; Isabella & Belsky, 1991). Our approach affords a holistic perspective on when and how mother and infant coordinate their neural, physiological, and behavioral responses. The results revealed that 4- to 6-month-old infants and their caregivers show INS associated with proximity and social touch. The link between touch and INS provides crucial new insights on the aspects of interaction quality that support caregiver-infant neural alignment. These findings pave the way for further research into the role of INS in facilitating the social bond between caregiver and child as well as the promotion of children’s social learning (Atzil et al., 2018; Feldman, 2017). The relation between touch and INS has implications for clinical interventions comprising social touch, such as kangaroo care, and could shed light onto a potential neural mechanism for why close contact is beneficial for infant development (Hardin et al., 2020). Synchronization in close physical contact could also promote children’s social learning just as it is suggested during social interactions (Piazza et al., 2021; Wass et al., 2021). It would be important to examine further outcome variables of learning during close contact and whether this relation is mediated by INS. IPS, on the other hand, was related to dyads’ co-regulation when infants expressed negative affect. These first results on the different levels of synchrony pave the way toward the future examination of the coupling of brain activation and RSA, especially in infancy, to provide a deeper understanding of the link between body and brain (Beauchaine, 2015). Taken together, the present study highlights the exciting opportunities of multi-level hyperscanning to uncover neurobiological pathways of early social communication and to provide a deeper understanding of the link between body and brain in human development.

## 5. Acknowledgments

We are grateful to Gabriela Markova and Sam Wass for their advice as well as our student assistants Lucie Zimmer, Katharina Hager and Marlies Schermann for their assistance in data acquisition, and video coding. Additionally, we thank all families who participated in the study and the Department of Obstetrics and Gynecology of the Vienna General Hospital for supporting our participant recruitment.

## CRediT author statement

**Trinh Nguyen**: Conceptualization, Methodology, Software, Formal analysis, Investigation, Data Curation, Project administration, Writing - Original Draft, Visualization. **Drew Abney:** Methodology, Software, Writing - Review & Editing. **Dina Salamander**: Investigation, Data Curation, Project administration. **Bennett I. Bertenthal:** Methodology, Writing - Review & Editing. **Stefanie Hoehl**: Conceptualization, Methodology, Resources, Writing - Original Draft, Supervision, Funding acquisition.

## Open Practices

The study was formally preregistered on https://aspredicted.org/blind.php?x=6hy5n6. The preprint was made available at biorxiv (https://doi.org/10.1101/2021.01.21.427664) under a CC-BY 4.0 International license. fNIRS and RSA data as well as MATLAB and RStudio analysis code will be made publicly accessible on OSF upon publication. The conditions of our ethics approval do not permit public archiving of video data. Readers seeking access to the data should contact the lead author Trinh Nguyen. Access will be granted to named individuals in accordance with ethical procedures governing the reuse of sensitive data. Specifically, requestors must meet the following conditions to obtain the data: completion of a formal data sharing agreement.

