## Supplementary Information for "Proximity and Touch are associated with Neural but not Physiological Synchrony in Naturalistic Mother-Infant Interactions"

S1: fNIRS analyses

Control Analyses for INS

We compared original coherence values against random pair coherence values for each dyad and channel combination to differentiate significantly increased neural synchronization from spurious correlations. Generalized linear mixed models (GLMM) included wavelet transform coherence (WTC) values of changes in oxygenated hemoglobin (HbO) as the response variable and the fixed and interaction effects of pairing (original vs. random), condition (distal watching vs. proximal watching vs. free play) and region of interest (inferior frontal gyrus [IFG] vs. lateral prefrontal cortex [lPFC] vs. medial prefrontal cortex [mPFC]). We assumed a random slope for all fixed effects in random intercepts for each dyad.

The model revealed a significant fixed effect of pairing, $\chi^2(1)=114.94$, $p<.001$, condition, $\chi^2(2)=38.31$, $p<.001$, region of interest, $\chi^2(2)= 29.37$, $p<.001$. In addition, all interaction effects between pairing and condition, $\chi^2(2)=21.08$, $p<.001$, pairing and region of interest, $\chi^2(2)=88.34$, $p<.001$, condition and region of interest, $\chi^2(4)=13.55$, $p=.009$, as well as the three-way interaction between pairing, condition, and region of interest, $\chi^2(4)=24.02$, $p<.001$. In post-hoc analyses, we contrasted original and random pairing in each region of interest for each condition (FDR-corrected). Synchronization during the distal watching condition was significantly higher in original dyads in comparison to random dyads in the IFG, $t=7.18$, $p<.001$, but in the lPFC or mPFC, $p>.090$. The proximal watching condition showed increased INS in the IFG and lPFC in original vs. random pairings, $t=3.88\text{-}8.23$, $p<=.001$. The free play condition showed significant INS in all regions of interest, $t=3.79\text{-}8.00$, $p<=.002$.

Additional INS analysis

To control for confounds in coherence values due to length of the epochs, we also analysed the difference in the two non-interactive and one interactive condition by splitting the latter into 90-second epochs. Coherence values, i.e., INS were entered as the response variable. Fixed effects included condition (distal watching vs. proximate watching vs. free play 1 (1-90s) vs. free play 2 (91-

180s) vs. free play 3 (181-270s)) and regions of interest (IFG vs. lPFC vs. mPFC) as well as their interaction effect. We assumed a random slope for all fixed effects in random intercepts for each dyad.

The GLMM output revealed a significant fixed effect of condition, $\chi^2(4)=33.00$, $p<.001$, region of interest, $\chi^2(2)=20.41$, $p<.001$, as well as their interaction, $\chi^2(8)=27.57$, $p<.001$. Post-hoc contrasts between conditions in the regions of interest showed that the conditions did not differ from one another in the IFG, $p>.449$. The proximal watching and the free play conditions showed higher INS than in the distal watching condition, $t>3.86$, $p=0.001$, but were not different from one another in the lPFC, $p>.053$. All epochs of the free play conditions showed higher INS in the mPFC in comparison to the distal watching condition, $t>3.00$, $p<.022$. The third epoch from the free play condition also showed higher INS than in the proximal watching condition, $t=3.19$, $p=.013$. There were no further significant differences in INS between conditions, $p>.052$.

Taken together, the differences in INS between the free play condition in comparison to the distal watching condition remain the same. The contrast in INS between the proximal condition and the free play condition are significant in the third epoch of the free play condition. This result indicates that INS increased over time during the free play condition, even though the contrasts between free play epochs were not significant. Still, we rule out the effect of measurement duration on INS.

HbO INS cross-correlation analysis

Next, we explored condition-related differences when we use cross-correlation instead of wavelet-transform coherence to estimate INS. 0-lag cross-correlation values were Fisher's z transformed and included in the linear mixed effects model as the response variable. Again, we included fixed effects for condition, region of interest and their interaction effect. A full random effects structure including random slopes for condition, region of interest and the interaction between both variables as well as random intercepts for each dyad was modelled. The results showed that there were no significant differences between conditions, region of interest, $p>.355$. Moreover, the interaction effect was significant as well, $p=.780$.

HbR INS analysis

We also ran the control analyses for neural synchrony in deoxygenated hemoglobin changes (HbR). We were able to replicate the results from HbO WTC analysis, but differences between original and random pairs were even stronger, specifically depicted in the main effect of pairing, $\chi2(1)=378.97$, $p<.001$. In post-hoc contrasts, we find that original pairs showed higher INS in all conditions and all regions of interest in comparison to random pairs, $t>11.80$, $p<.001$.

We analyzed the differences between conditions for HbR neural synchrony. The GLMM included HbR synchrony values as the response variable with fixed and interaction effects of condition and region of interest. A full random effects structure was assumed including random slopes for all fixed and interactions effects as well as random intercepts for each dyad.The findings show that fixed effects for condition, $\chi^2(2)=14.89$, $p<.001$, and region, $\chi^2(4)=48.97$, $p<.001$, as well as their interaction was significant, $\chi^2(8)=18.12$, $p=.020$. The findings for HbO and HbR differed somewhat when we subsequently compared the conditions in each region. Significantly increased neural synchrony during Free Play in comparison to distal watching was detected in bilateral lPFC and mPFC, $t>2.37$, $p<.046$. The proximal watching condition showed higher neural synchrony than the distal watching condition in the lPFC, $t=2.63$, $p=0.022$. The free play condition and proximal watching condition did not differ in synchronization in the mPFC, $p>.136$. None of the conditions differed in neural synchrony in bilateral IFG, $p>.779$.

HbR INS cross-correlation analysis

Here, we explored condition-related differences of 0-lag cross-correlation scores of HbR time-series from mothers and infants. 0-lag cross-correlation values were Fisher's z transformed and included in the linear mixed effects model as the response variable. Again, we included fixed effects for condition, region of interest and their interaction effect. A full random effects structure including random slopes for condition, region of interest and the interaction between both variables as well as random intercepts for each dyad was modelled. Similar to HbO INS cross-correlation analysis, the results showed that there were no significant differences between conditions, region of interest, p>.262. Moreover, the interaction effect was significant as well, p=.691.

Infant Touch and HbR INS

Next, HbR WTC values from the Free Play condition were tested as the response variable. The findings revealed a significant fixed effect of stimulating touch durations, $\chi^2(1)=4.52$, $p=.033$. Longer durations of stimulating touch were again related to lower INS, estimate=-0.067, SE=0.031, 95% CI=[-0.129 -0.005]. All other touch durations were not significantly related to INS, $p>.173$.

Infant Affect and INS

Next, we assessed infant affect as a behavioral correlate of INS. Again, HbO WTC values from the Free Play condition were included as the response variable. Infant positive and negative affect durations were however separately tested as fixed effect variables due to collinearity. The issue was caused by an expected high correlation between durations of infant positive and negative affect, $\beta=-0.477$, $p=0.001$. Still, neither negative nor positive affect duration was significantly related to INS, $p=.404$ (FDR-corrected).

S2: RSA analyses

Control analyses for Physiological synchrony

To detect significant and meaningful synchronization in RSA, we conducted a control analysis using shuffled time series (Abney et al., 2021). We compared original and shuffled pairs (fixed effect) in %DETerminism (%DET) in each condition (fixed effect). We assumed random intercepts for each dyad. The findings show that original pairs showed significantly higher values in %DET in all conditions in comparison to shuffled pairs, indicated by the highly significant fixed effect of pairing, $\chi^2(1) > 244.38$, $p < .001$. All post-hoc contrasts reveal higher values for original pairs than shuffled pairs, $t > 10.77$, $p < .001$.

Physiological synchrony assessed by cross-correlation

We also examined whether mother and infant showed physiological synchronization by investigating the lag-0 cross-correlation of their changes in RSA over time. Findings also revealed that mother-infant dyads did not generally covary in their changes of RSA throughout the distal watching, the proximal watching or the free play condition as evidenced by a non-significant comparison with their shuffled counterpart, $p > .442$. The original pairs also did not show significant differences between the conditions, $p = .378$.

Infant affect and physiological synchrony

The second analysis looked at correlations between IPS and infant affect. We had to calculate two linear regressions due to collinearity. Both regressions comprised %DET as the outcome variable. Infant positive affect duration was first assessed as a predictor variable. Here, the results show that higher durations of infant positive affect were related to lower IPS, $\beta = -0.390$, $SE = 0.142$, *95% CI*=[-0.677 -0.104], $t = -2.74$, $p = .009$ (FDR-corrected). A second regression assessed the correlation between IPS and negative infant affect duration and revealed a strong positive correlation, $\beta = 0.538$, $SE = 0.130$, *95% CI*=[0.275 0.800], $t = 4.14$, $p < .001$ (FDR-corrected). The results are depicted in *Figure S2*.

S3: Behavioral analysis

Infants' gaze during distal and proximal watching conditions

To assess and control for attentional processes in the non-interaction watching condition, we compared infants' gaze directions towards the screen, the mother's face, and away in the distal and proximal watching conditions. We used a linear regression to test for differences in condition and direction. The findings depict a significant effect for direction, $\chi^2(2) = 813.30$, $p < .001$, and the interaction between direction and condition, $\chi^2(2) = 6.49$, $p = .002$. The was no significant difference in gaze durations between conditions, *p*=.449. Post-hoc analyses controlled for multiple comparisons showed that infants' gaze towards the screen did not differ in duration between conditions (distal*:* *estimate*=72.45, *SE*=1.83, *95% CI*=[68.86 76.06]; proximal: *estimate*=72.47, *SE*=1.83, *95% CI*=[68.87 76.07]). Gaze directed towards the mother was significantly longer in the distal watching condition, *estimate*=8.80, *SE*=1.83, *95% CI*=[5.20 12.40], than in the proximal watching condition, *estimate*=-0.57, *SE*=1.83, *95% CI*=[0.00 4.17]. Gaze directed away was marginally longer in the proximal watching condition, *estimate*=15.57, *SE*=1.83, *95% CI*=[11.97 19.17], than in the distal watching condition, *estimate*=10.75, *SE*=1.83, *95% CI*=[7.16 14.36].

Supplementary Tables

Table S1. Description of categories used to code infant gaze during the non-interactive watching conditions and maternal touching behavior and infant facial affect during the interactive free play condition.

| Coding category | Description | Interrater-Reliability (Kappa) |
|---|---|---|
| **Non-interactive Watching Conditions** | | |
| **Infant Gaze** | | 0.91 |
| Gaze to screen | Gaze directed towards the tablet | 0.94 |
| Gaze at mother | Gaze directed towards the mother’s face | 0.92 |
| Gaze away | Gaze directed away from the tablet and the mother’s face, directed towards the room etc. | 0.88 |
| **Interactive Free Play Condition** | | |
| **Maternal Touch** | | 0.79 |
| Active Touch | Dynamic and stimulating forms of touch | 0.79 |
| Affectionate Touch | Stroking, massaging, and other gentle movements, caress/rub, and pat/tap, kiss | 1.00 |
| Stimulating Touch | Tickling, lifting, and rhythmic touch, playful and engaging touch such as squeeze/pinch/grasp, tickle/finger walk/poke/prod/push, shake/wiggle, and pull/lift/extension/clap | 1.00 |
| Passive Touch | Passive or static forms of touch, e.g. resting hands on infant’s body, passive bodily contact | 0.71 |
| Functional Touch | Touches used with instrumental intention, e.g. adjusting infant’s clothing, wiping mouth, removing cables | 0.73 |

| | | |
|---|---|---|
| No Touch | No physical contact between mother and infant | 0.89 |
| **Infant facial affect** | | 0.82 |
| Positive Affect | Smiling with mouth turned upward (open or closed), contraction of cheek muscle and/or under-eye muscle | 0.81 |
| Neutral Affect | Neutral facial expression | 0.80 |
| Negative Affect | Expressing negative emotions: distress, fretting, anger, discontentment, sadness or “pout face” as indexed by narrowed eyes, mouth curled or grimacing, lowered brows, mouth corners turned down | 0.86 |

Supplementary Figures

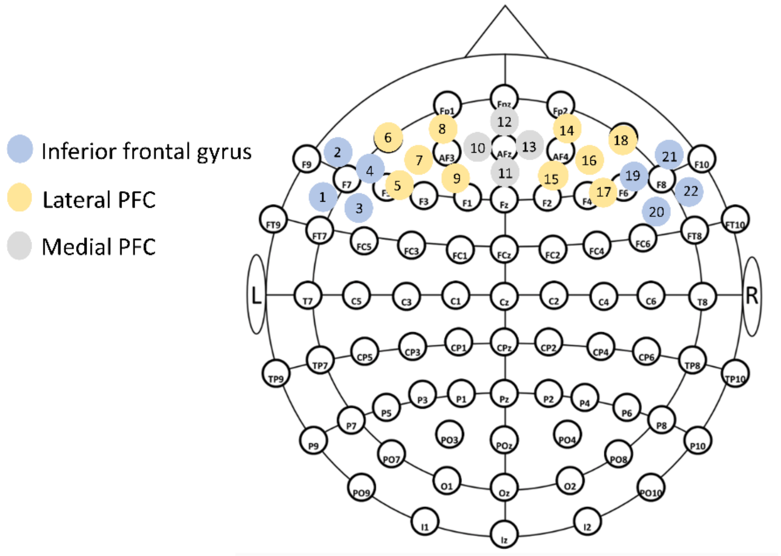


*Figure S1.* The channel configuration of the mother and infant cap comprises the following brain areas: Inferior frontal gyrus (1-4; 19-22), lateral prefrontal cortex (PFC; 5-9; 14-18), and medial prefrontal cortex (10-13).

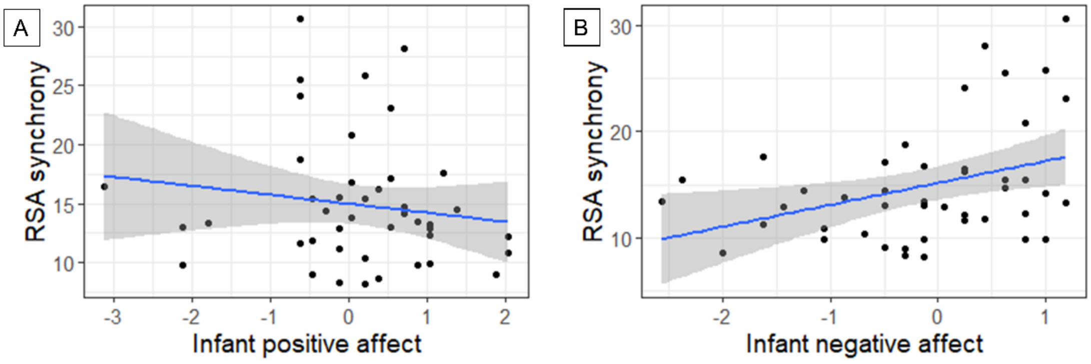


*Figure S2*. Plots depict (A) the negative correlation between duration of infant positive affect (x-Axis) and (B) the negative correlation between infant negative affect (x-Axis) and physiological synchrony (y-Axis) during the free play condition.
